# The High Transmission of SARS-CoV-2 Omicron (B.1.1.529) Variant is Not Only Due to Its hACE2 binding: A Free Energy of Perturbation Study

**DOI:** 10.1101/2021.12.04.471246

**Authors:** Filip Fratev

## Abstract

The mutations in the spike protein of SARS-CoV-2 Omicron variant (B.1.1.529 lineage) gave rise to questions, but the data on the mechanism of action at the molecular level is limited. In this study, we present the Free energy of perturbation (FEP) data about the RBD-hACE2 binding of this new variant.

We identified two groups of mutations located close to the most contributing substitutions Q498R and Q493R, which altered significantly the RBD-hACE2 interactions. The Q498R, Y505H and G496S mutations, in addition to N501Y, highly increased the binding to hACE2. They enhanced the binding by 98, 14 and 13 folds, respectively, which transforms the S1-RBD to a picomolar binder. However, in contrast to the case in mice the Q493R/K mutations, in a combination with K417N and T478K, dramatically reduced the S1 RBD binding by over 100 folds. The N440K, G446S and T478K substitutions had lesser contribution. Thus, the total effect of these nine mutations located on the interaction surface of RBD-hACE2 turns out to be similar to that observed in the Alpha variant. In a special circumstances it could be further altered by the E484A and S477N mutations and even lower binding capacity is likely to be detected. Finally, we provide a structural basis of the observed changes in the interactions.

These data may explain only partially the observed in South Africa extremely high Omicron spread and is in support to the hypothesis for multiple mechanisms of actions involved in the transmission.

**Graphical abstract:** 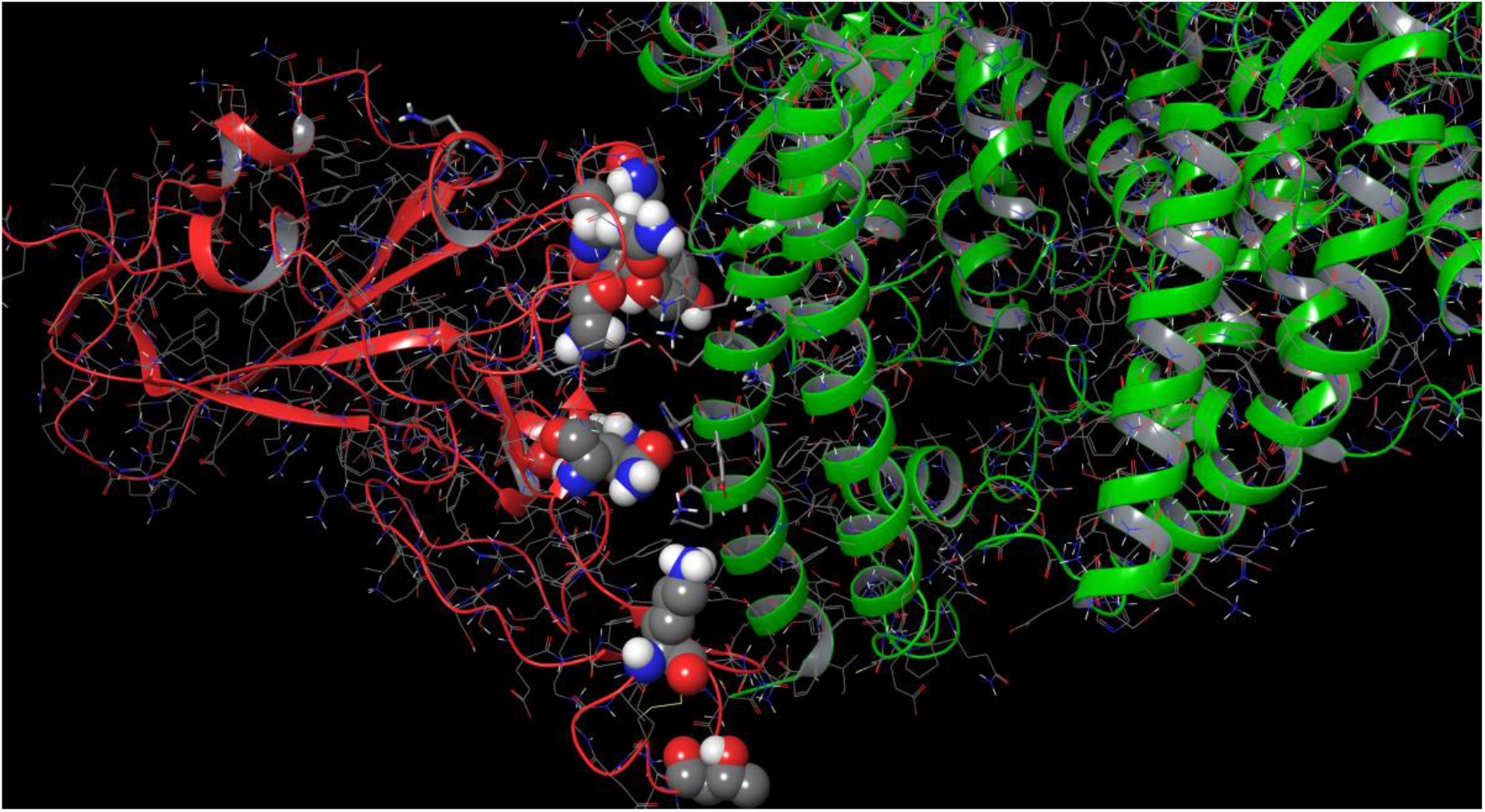

## Introduction

On November 26, 2021, WHO identified the SARS-CoV-2 Omicron variant (B.1.1.529 lineage) as a variant of concern (VOC), based on evidence that it contains numerous mutations that may influence its behavior [1]. However, the mode of transmission and severity of the Omicron variant remains unknown. The B.1.1.529 lineage has a total number of 11 amino acid mutations in its receptor binding domain (RBD): K417N, N440K, G446S, S477N, T478K, E484A, Q493R, G496S, Q498R, N501Y and Y505H [2]. These mutations have been discovered in samples collected in Botswana on November 11, 2021 and South Africa on November 14, 2021. As of 3 December and since 2 December 2021, 30 additional SARS-CoV-2 Omicron VOC cases have been confirmed in the European Union (EU). Countries and territories outside the EU have reported 377 confirmed cases. Globally there were 486 confirmed cases reported by 35 countries. It is uncertain if the Omicron variant is more transmissible or severe than the Delta variant. However, this is the most divergent strain detected so far during the pandemic raising concerns and several countries have already closed their borders due to the possible risk. The mutations in the spike protein of Omicron gave rise to questions, but the data on the mechanism of action at the molecular is level is limited.

In the end of December 2020 we provided urgently high quality data about the effect of the N501Y and K417N mutations, detected in Alpha and Beta variants, to the RBD-hACE2 binding by the Free energy of binding (FEP) approach [3]. The results have been later confirmed by the experimental studies [4-7]. Further, we performed a similar study also for the R346K mutation found in the *Mu* SARS-CoV-2 variant, which affected monoclonal antibodies (mAbs) from class 2 [8]. In the current study we used the same technique to assay the binding of the Omicron’s S1 RBD to hACE2 and to provide at a molecular level a detail description of the RBD-hACE2 interactions. The FEP method is one of the most successful and precise *in silico* techniques for protein-protein interactions predictions [9]. It outperforms significantly the traditional molecular dynamics based methodologies, such as for example MM/GBSA and empirical solutions like FoldX. It also often precisely predicts the free energy differences between the mutations [10-12] and has a more than 90% success in the prediction whether one mutation will have either a negative or positive effect on the binding [9].

## Results and Discussion

Our FEP calculations were based on the recently published X-ray structures of the Alpha and Beta SARS-CoV-2 RBD variants of the virus [13]. We observed that two groups of Omicron mutations significantly altered but with an opposite effect the S1 RBD-hACE2 binding (**Table 1, Figure 1** and **Figures 2A** and **2B**).

**Table 1.** FEP results for the selected mutations (kcal/mol) in Alpha, and Beta. The experimental values, if available, are also shown. See the text for details. Mode/Ref note whether the mutation increase (I) or decrease (D) the binding and reference, respectively. N/A – not available

| Mutation | $\Delta\Delta G$ (FEP)<br>in Alpha | $\Delta\Delta G$ (FEP)<br>in Beta | $\Delta\Delta G$ (exp) | Mode<br>/Ref |
| --- | --- | --- | --- | --- |
| Q498R | -1.92 | -1.98 | N/A | I |
| G496S | 0.04 | -0.23 | N/A | I |
| Y505H | -0.17 | -0.86 | N/A | I |
| Q493R | 2.01 | 2.43 | N/A | D |
| Q493K | 2.79 | 3.19 | N/A | D |
| G484A | 0.04 | - | N/A | D |
| N440K | -0.20 | - | N/A | I |
| G446S | 0.21 | - | N/A | D |
| K417N | 0.43 | - | 0.96 | D<sup>14</sup> |
| S477N | -1.18 | - | -0.33÷-0.72 | I<sup>14</sup> |
| T478K | 0.16 | - | - | D |
| N501Y | 1.46 | - | 1.43 | I<sup>3,14</sup> |

**Figure 1.**
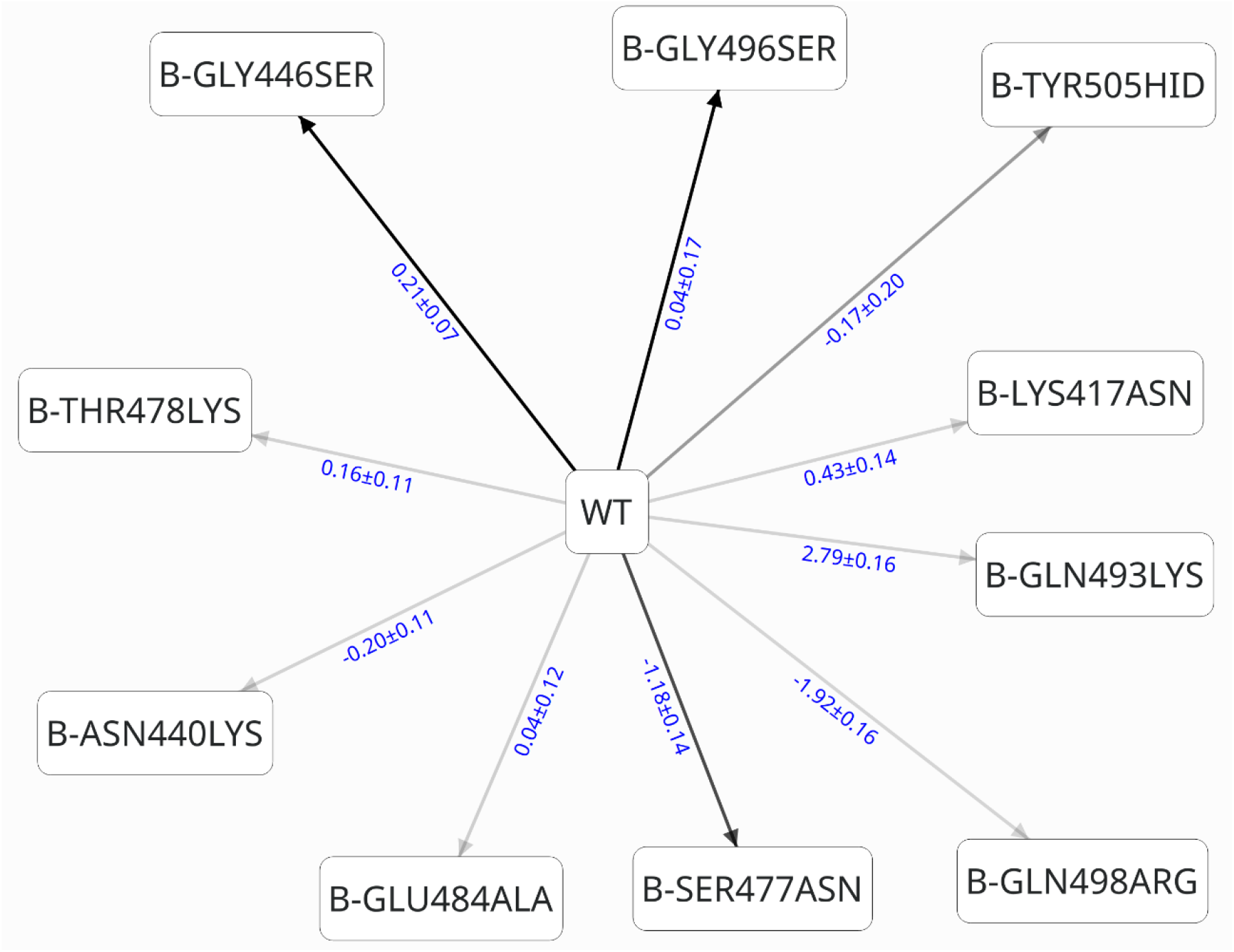
A schematic presentation of the perturbation networks and calculated free energies of binding using the structure of Alpha variant of SARS-CoV-2. In blue color are shown the calculated by Bennett approach ΔΔG values, respectively.

**Figure 2.**
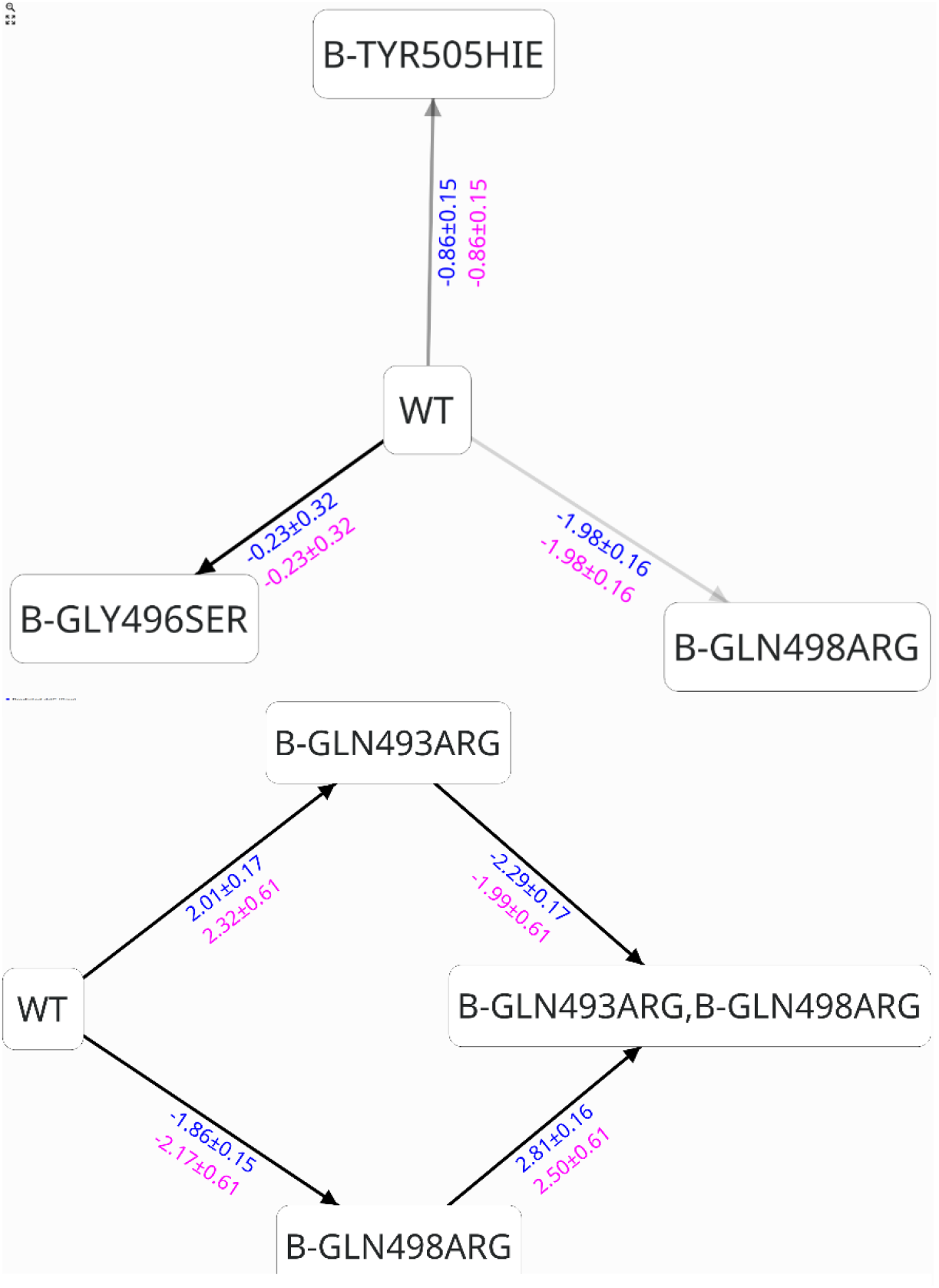
A schematic presentation of the perturbation networks and calculated free energies of binding using (***A***) the structure of Beta and (***B***) Alpha variants of SARS-CoV-2. In blue and red color are shown the calculated by Bennett and cycle closure approach ΔΔG values, respectively.

### The impact of Q498R and surrounding mutations which contributes to the higher RBD-hACE2 binding

The Q498R substitution was in the center of the constellation of mutations which contributes to the higher binding and showed the most significant change in the free energy: **-1.92 kcal/mol**. This value was obtained based on the Alpha variant structure but was also further confirmed by an independent FEP calculation in the Beta variant (−1.98 kcal/mol). To this group belong also the Gly496Ser and Tyr505His mutations. In the Beta variant we calculated ΔΔG values of -0.23 kcal/mol and -0.86 kcal/mol, respectively. However, in the

Alpha variant they showed a free energy of binding change of zero and -0.17 kcal/mol, respectively. We analyzed this difference and concluded that the conformation of Asn498 is an important factor for the observed variations and in particular the carbonyl group, which can have opposite conformations. In the employed experimental structures this was also seen and both conformations have been resolved [13]. Hence, to track the possible influence of this phenomena we used one of this conformations in our Alpha model and one in our Beta model, before to start the calculations, which produced these deviations in the FEP values. We observed that when the carbonyl group is with an orientation toward His505, as in our Beta model, the Ser496 and His505 contributed significantly higher to the binding. To confirm this hypothesis we changed the conformation of Gln498 in our initial Alpha model to those seen in the Beta variant and was already employed for abovementioned calculations. As a result we obtained a free energy difference for G496S and Y505H of -0.35 kcal/mol and -0.34 kcal/mol, respectively, which confirms our hypothesis about the conformation of Gln498. The medium values for based on all these calculations can be estimated to be in a ΔΔG range of **-0.2÷0.3 kcal/mol** and **-0.4÷0.5 kcal/mol** for G496S and Y505H mutations, respectively.

Thus, if we consider that the N501Y mutation was present in both Alpha and Beta variants and the available experimental data we can assess the RBD-hACE2 binding of discussed above substitutions in K_d_ values by the roughly estimating formula ΔG=RTlnK_d_. Based on this we calculated that the Q498R, Y505H and G496S mutations, in addition to N501Y, highly increased the binding to hACE2. They enhanced the binding by 98, 14 and 13 folds, respectively, which transforms the S1-RBD to a picomolar binder. The RBD binding to hACE2 receptor for Alpha and Beta variants has been experimentally assessed to have K_d_ values of 8.1nM and 13.3nM, respectively [14]. Only the Q498R mutation in Omicron decreases the K_d_ value of the RBD-hACE2 binding to 0.27nM whereas the measured in Beta variant contribution of Y505H mutant provide an K_d_ value of 1.9nM. A couple less contributing residues in this hot spot region were also detected. The N440K and G446S had a small effect to the RBD-hACE2 binding and the ΔΔG values were -**0.20** and **0.21 kcal/mol**, respectively; i.e. in frame of possible error. We speculated that the G446S in a combination with G498R and surrounding residues may additionally stabilize this constellation of harmful mutations and in particular Arg498 but this turn out to be not true. The calculated ΔΔG value based on our FEP generated Omicron averaged structure for this portion of the S1 RBD was 0.24 kcal/mol.

### The impact of Q493R and surrounding mutations which contributes to the lower RBD-hACE2 binding

Surprisingly, we observed a high negative impact to the binding of Q493R mutation and even higher for the possible Q493K substitution. We expected some decrease, similar to the case of K417N, due to the close positions of other positively charged residues of hACE2 such as Lys31 (see below), but no so significant impact. Moreover, it seems that the general belief at the moment is that Q396R/K mutation contributes, like N501Y, for an increases of RBD-hACE2 binding, which is based mainly on the mouse *in vitro* and *in vivo* models [15-16].

However, it is probably less known that the K417N in mice also acts in a different way than in human increasing the binding [16]. Initially, for the Q493K mutation we obtained a ΔΔG value of 2.79 kcal/mol whereas for the Q493R substitution a ΔΔG value equal to 2.21 kcal/mol, using the Alpha variant of the structure as a base for our calculations. In Beta variant RBD-hACE2 complex the values were similar and even higher (ΔΔG=3.19 and 2.43 kcal/mol). Initially, we thought that this is some error due to the possible bias of the transformation of neutral to charged residues observed often in FEP calculations [17]. However, it was strange that we observed that during the comparison of two identical substitutions of Gln to Arg substitutions at positions 493 and 498, respectively. Further, we did the reverse mutations, i.e. Arg493 to Asn and obtained a similar ΔΔG value values with opposite sign: R493Q had ΔΔG value of -2.31 kcal/mol.

Next, we started to discuss some structure based related explanation, which eventually can provide also a molecular basis of the observed Q493R negative contribution to the formation of the RBD-hACE2 complex. Looking on this complex one can see some important differences of the interacting residues close to Q493R in humans and mice (**Figures 3A** and **3B**). In particular, this was the Lys43 to Asn mutation in mice ACE2 (mACE2). Thus, it seems that in this hot spot of interactions there is a significant difference in mice and humans and the K37N substitution in ACE2 presumably plays a major role in the interspecies difference. To validate this hypothesis we studied how the RBD_MASCp36_ variant, which contains the N501Y, Q493H and K417 mutations, binds to mACE2. The structure of RBD_MASCp36_-mACE2 complex has been recently obtained and we mutated the His493 to its native Glu394 residue [16]. As a results, we obtained a ΔΔG value of 1.07 ±0.17 kcal/mol. Also, we calculated the impact of H493K mutation and obtained a further increase of the binding of -0.62 ±0.17 kcal/mol whereas the Q493H in RBD-hACE2 provided a stabilization of -0.2 kcal/mol, based on the formation of H-bonds and ionic interactions with Lys31 and Glu35 in hACE2. These data are in an excellent agreement with the experimental studies demonstrating that the FEP calculations correctly described the RBD-mACE2 interactions in this part of the binding surface. In addition, the experimental data showed that the K417N mutation enhanced the RBD binding to mACE2 from K_d_=12.7µM to K_d_ value of 2.4µM whereas in hACE2 the Lys417 decreased it from K_d_=12.7nM to K_d_ 19.9nM [16]. These data clearly demonstrate that our hypothesis about the interspecies difference of RBD-ACE2 interactions in this hot spot of mutations is correct and the calculated by FEP impact of Q493R mutation is correct.

**Figure 3.**
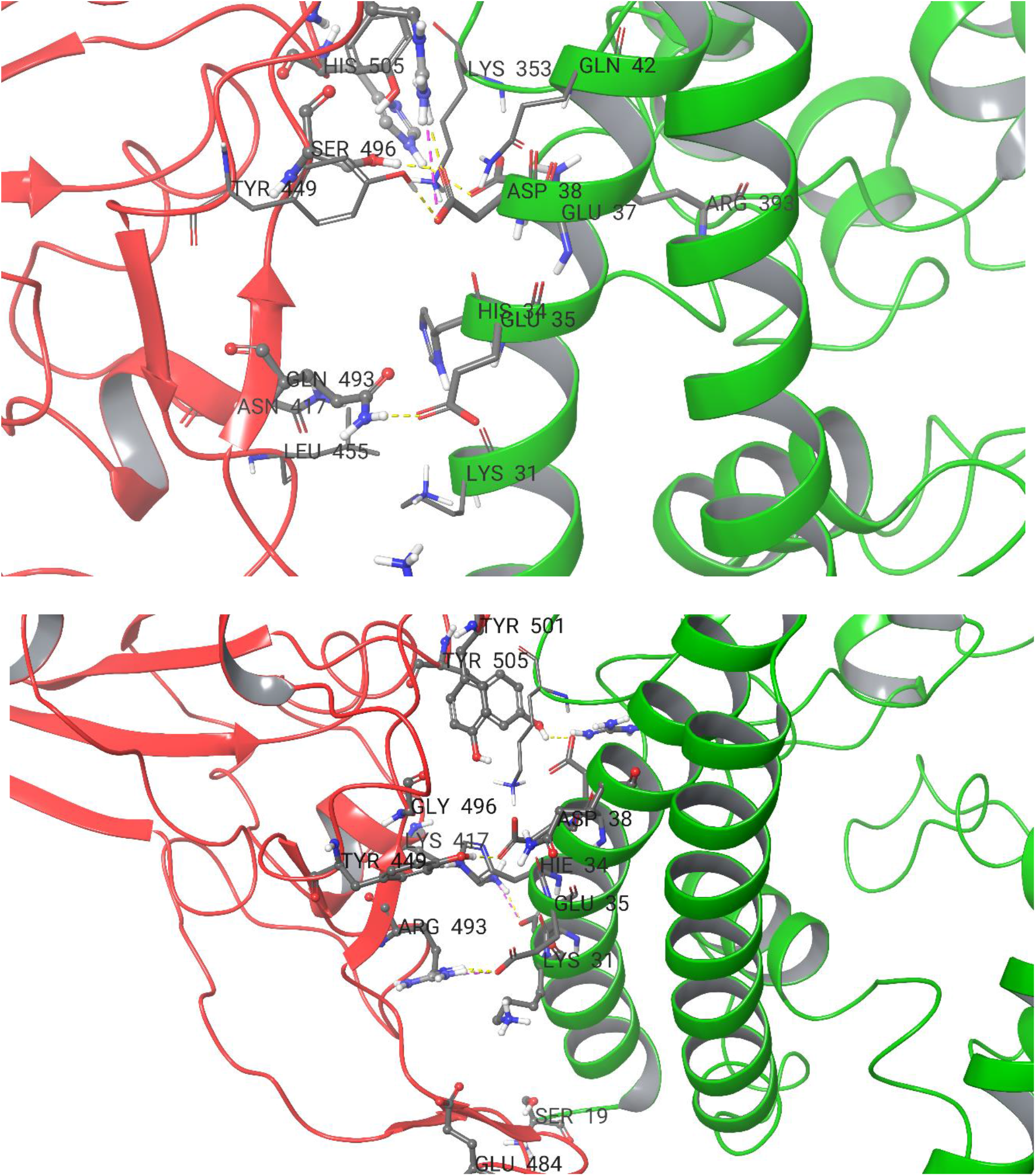
A close look of the identified by FEP interactions between the S1 RBD of ***(A)*** mutations close to Q498R and ***(B)*** Q493R substitutions, respectively and hACE2. With yellow dot lines are shown the H-bonds. With green and red colure are shown the hACE2 and the Omicron’s RBD binding surfaces.

### The common effect of all detected in the RBD of Omicron mutations

Finally, we summarized the obtained data obtained for the detected two hot spots of so many mutations in the RBD of Omicron. In order to have a more precise value of the effect of Q493R and Q498R mutations we performed additional cycle closure FEP calculations (**Figure 2B**) and obtained ΔΔG values of **2.32±0.61** kcal/mol **-2.17±0.61** kcal/mol, respectively. *Based on all these data one can conclude that the total effect of Q493R and Q498R mutations lead to almost no change in the binding of the RBD to hACE2*. Further, we have a constellation of other mutations, which we described herein or previously and for some of these mutations there are also solid experimental data. Based on this information one can conclude also that *the common effect of K417N, G496S and Y505H mutations did not also provided a significant total change*: **0.96** kcal/mol experimental value for K417N *vs* about -**0.6÷0.8** kcal/mol for G496S and Y505H calculated here. The N440K, G446S and T478K substitutions had lesser contribution and mutually effect excluded too. Thus, we concluded that the Omicron’s RBD-hACE2 is about those observed in the Alpha variant.

However, it may also be a little further affected in two opposite directions. The eventual increase of the binding can be contributed by the S477N and T478K mutations. The calculated by FEP effect to the S1 RBD-hACE2 binding of E484A has no any impact. However, it can indirectly influence the effect of Q493R mutation and may explain the observed differences between calculated free energy values in Alpha and Beta (see below).

### Comparison to the available experimental data

Our data correspond well to the already available experimental assays for some of the studied mutations and a comparison is listed in **Table 1**. However, we need also to consider two main factors before to make a general conclusions. Indeed, between the various experimental SARS-CoV-2 RBD structures there are differences in terms of resolution, quality, experimental approach used, conformation of specific residues etc. and we can expect some slight deviations from the experimental data. The same is valid also for our previous calculations [7-8]. For instance, for the N501Y mutation in our FEP study we obtained a ΔΔG values of 1.54 kcal/mol, which is identical to the experimental results [14]. For the K417N we obtained previously a ΔΔG value of +0.5 kcal/mol and 0.43 kcal/mol herein. However, during our calculations for the *Mu* variant of the virus we also obtained a value of 1.29 ± 0.44 kcal/mol. Even though these values are close to the experimentally observed value of +0.68 kcal/mol [14] we can see some slight deviations but they are in a frame of both the experimental and FEP calculation error. The other major factors in FEP which may produce errors are the bad sampling and convergence [11]. The convergence of all mutations described herein was good but for more precise data an addition increase of both the simulation length and also cycle closure calculations can be performed.

Note also that the impact of S477N mutation is overestimated due to the artificially introduced in this study cap of the last resolved ACE2 residue Ser19. The latter interacts and provides better binding with Asn477. In the real case this could be different but to make a model of the ACE2 part for a FEP study is not practical at this point of the study and can introduce even more non-native interactions. However, the sign of the calculated ΔΔG is correct and the value is not so different than the experimental assays which showed ΔΔG= -0.33, -0.67 and -0.72 kcal/mol [14].

### Molecular basis of the Omicron’s S1 RBD-hACE2 biding and observed changes in the free energies

A brief description of molecular basis of the observed differences in the binding and interactions between the S1 RBD of Omicron and hACE2 can be provided (**Figure 3A** and **3B**). The Q498R, Y505H and G496S altered significantly the RBD-hACE2 interactions in both the Alpha and Beta variants. The Arg498 provides better stabilization of RBD-hACE2 complex by electrostatic interactions and H-bond with Glu37 of hACE2 but also provide stabilization to Tyr501. The Ser496 is also involved in these interactions which makes an H-bond with Glu37. The His505 keeps the H-Bond with Asp38 and alongside with Tyr501 and Arg498 creates a bridge of stable π-π stacking and ionic interactions in this hot spot of mutations.

Instead, the Q493R has unfavorable interactions with Lys31 of hACE2 which destabilizes the binding in this part of the RBD-hACE2 complex and repulse the Arg493 from the binding surface, although an H-bond with Glu35 can be seen in the averaged structure (**Figure 3B**). Here, we can have different scenarios in the SARS-CoV-2 variants. In Alpha variant we observed a rapid movement of Arg493 toward the Glu384 and long-range electrostatic interactions between them. In Beta variant the E484K substitution is present which additionally destabilize the RBD-hACE2 complex. In simple words the Arg493 cannot adopt a stable conformation in both the pocket occupied by Lys484 and those formed between Lys31, His34 and Glu37 from hACE2 due to the unfavorable electrostatic interactions and its big size, respectively. In addition, the Leu455 also acts as a barrier. This explains well observed by our FEP calculations additional decrease of the RBD-hACE2 binding in Beta variant. We speculate that the E484A mutation will play a significant role for the Arg493 flexibility and conformational changes and may have an indirect impact to the Omicron binding to hACE2.

The difference between Gln493Arg and Gln493Lys mutations come mainly from the size of these charged residues. We observed that Lys493 can approach to the ACE2’s pocket formed by Lys31, His34 and Glu37. It even may forms stable interactions with hACE2 in this region but in this way destabilizes the interactions formed by Arg498 and the other residues from this binding hot spot in Omicron. It also disturbs the interactions of Gln398 with hACE2 moving the residues a bit away from the binding surface. These can well explain the observed higher contribution of Lus493 than Arg493 to the decrease of the RBD-hACE2 binding.

### Some limitations and future work

The limitations of this study and the possible feature work which can bring some improvements are as follows:

- We performed only the default protocol of 5ns sampling which may be not enough in some cases as we previously shown [7]. However, in almost all cases the convergence was good and we did not expect any significant changes, only refrainment’s.
- The cycle closure FEP calculations for all perturbations will indeed bring an improvement. However, the urgency of the data provided and the time of the computations did not allowed us to do that.
- Additional studies of the interactions between individual mutations in Omicron variant and in particular the E484A contribution to the Arg493 destabilization should be performed.
- A deeper study about the possible impact of the eventual bias produced by the perturbations of neutral to charged residues may be performed but we believe that this is out of the scope of this study.

## Methods

The methods used have been described in details previously [7-12]. In short, to perform our FEP calculations and structural studies we used the recently deposited structures with pdb accession numbers 7EKF (for the Alpha variant of the RBD-hACE2 complex) and 7EKG (for the Alpha variant of the RBD-hACE2 complex), respectively. For RBD-mACE2 complex (SARS-COV-2 Spike RBDMACSp36 binding to mACE2) we used the pdb structure 7FDK. All protein preparations and calculations were performed by Schrödinger suite software [18]. To calculate the differences in the free energy of binding for each complex in this study we employed the Desmond FEP/REST approach described in details previously [7-12]. A sample scheme of the thermodynamical cycle for the calculations of the binding affinity change due to mutations in interacting protein-protein interface is shown on Figure 1 in ref [19]. The binding free energy change ΔΔG_AB_ or in simple ΔΔG can then be obtained from the difference between the free energy changes caused by the particular mutation in the bound state (ΔG_1_, complex leg) and the unbound state i.e. in solvent (ΔG_2_, solvent leg). In a typical FEP calculation for a mutation from state A to state B, several perturbation lambda (λ) windows are needed in order to obtain a smooth transition from the initial state A to the final state B. The default sampling FEP+ protocol was applied here with the number of λ windows either 12 or 24, in dependence of the mutation charge; i.e. same or different. An equilibration and 5 ns-long replica exchange solute tempering (REST) simulations in a muVT ensemble was further conducted. Only the mutated atoms was included in the REST “hot region” region. OPLS4 force field was used for the all simulations [20]. The obtained average structures from the simulations were used for the structural comparison on **Figures 3A** and **3B**. All results can be reproduced and further analyzed by the Appending II files, which are available on request and include the structures, all parameters and all protocols, which are available on request.

## Acknowledgments

Thanks due to the Suman Sirimulla for the helpful discussion. We thanks to Dimitar Dimitrov which organized several activities for execution of this study and to Ivan Stankov for editing the manuscript.

## Data and Software Availability

All protein preparations and calculations were performed by Schrödinger suite 2021-2 software [22] https://www.schrodinger.com/downloads/releases]. All results can be reproduced and further analyzed by the provided on request Appending I necessary files, which include the structures, all parameters and protocols, respectively

